# Transcriptomic profiling of human orbital fat and differentiating orbital fibroblasts

**DOI:** 10.1101/2021.05.13.443857

**Authors:** Dong Won Kim, Kamil Taneja, Thanh Hoang, Clayton P. Santiago, Timothy J. McCulley, Shannath L. Merbs, Nicholas R. Mahoney, Seth Blackshaw, Fatemeh Rajaii

## Abstract

**Purpose:** Orbital fat hyperplasia has a central role in the manifestations of thyroid-associated orbitopathy (TAO). To better understand the pathways involved in adipogenesis in TAO, we have used transcriptomic methods to analyze gene expression in control and TAO patients, as well as in differentiating orbital fibroblasts (OFs).

**Methods:** We performed bulk RNA sequencing (RNA-Seq) on intraconal orbital fat to compare gene expression in control and TAO patients. We treated cultured OFs derived from TAO patients with media containing dexamethasone, insulin, rosiglitazone, and isobutylmethylxanthine (IBMX) to induce adipogenesis. We used single nuclear RNA-Seq (snRNA-Seq) profiling of treated OFs to compare gene expression over time in order to identify pathways that are involved in orbital adipogenesis *in vitro* and compared the dynamic patterns of gene expression identify differences in gene expression in control and TAO orbital fat.

**Results:** Orbital fat from TAO and control patients segregate with principal component analysis (PCA). Numerous signaling pathways are enriched in orbital fat isolated from TAO patients. SnRNA-Seq of orbital fibroblasts undergoing adipogenesis reveals differential expression of adipocyte-specific genes over the developmental time course. Furthermore, genes that are enriched in TAO orbital fat are also upregulated in orbital adipocytes that differentiate *in vitro*, while genes that are enriched in control orbital fat are enriched in orbital fibroblasts prior to differentiation.

**Conclusions:** Differentiating orbital fibroblasts serve as a model to study orbital fat hyperplasia seen in TAO. We demonstrate that the insulin-like growth factor-1 receptor (*IGF-1R*) and Wnt signaling pathways are differentially expressed early in orbital adipogenesis.

**Précis:** To understand the pathways involved in adipogenesis in TAO, we used transcriptomic methods to analyze gene expression in control and TAO patients, as well as in differentiating OFs. We demonstrate that the IGF-1R and Wnt signaling pathways are differentially expressed during orbital adipogenesis.

## Introduction

Thyroid-associated orbitopathy (TAO) is a form of autoimmune thyroid disease in which shared auto-antigens in the thyroid gland and orbit are targets of the immune response.^1–3^ About 25% of patients with autoimmune thyroid disease will develop the ocular disease, which can be vision-threatening. It is thought that inflammatory signaling causes expansion of the orbital soft tissues due to fibrosis and orbital fat hyperplasia which causes ocular morbidity due to corneal exposure and/or compressive optic neuropathy.^4–6^

The orbital soft tissue expansion that causes vision loss and other ocular morbidity associated with TAO is thought to be the final step of an inflammatory cascade that results in fibrosis and adipogenesis. Details of the mechanisms underlying orbital soft tissue fibrosis and adipogenesis are not well understood. Generally, adipogenesis is initiated by transcription factor cascades involving transient early expression of CCAAT/enhancer-binding protein (C/EBP) -β and -δ followed by induction of CEBP-α and peroxisome proliferator-activated receptor γ, transcription factors, which in turn induce expression of genes involved in terminal adipocyte differentiation.^7,8^ These pathways contribute to a generic adipocyte differentiation program. However, adipocyte metabolic function, morphology, preadipocyte proliferation and capacity for adipogenesis differ among adipose tissue depots.^9^ These molecular and physiological differences, which have been explored in subcutaneous, visceral, and brown adipose tissue depots, are thought to be intrinsic to adipocytes and may be related to their embryonic origins.^9^ The differences in orbital adipose tissue, and the way in which they may contribute to the TAO phenotype, however, have not yet been explored.

To identify molecular mechanisms controlling adipogenesis in TAO, we have conducted bulk RNA-Seq analysis of primary orbital fat from both control and TAO patients and also used snRNA-Seq to profile orbital fibroblasts undergoing adipogenesis *in vitro*.

## Methods

### Bulk RNA-Seq

Human intraconal orbital fat was obtained from TAO patients undergoing orbital decompression or controls undergoing routine resection of prolapsed orbital fat (Table 1), which has been shown to be intraconal fat,^10^ or enucleation using a protocol approved by the Johns Hopkins University Institutional Review Board and following the tenets of the Declaration of Helsinki. TAO patients were inactive, defined by a clinical activity score (CAS) less than 4.^11^ Samples were placed in RNA*later*™ (ThermoFisher Scientific) or on ice, depending on their subsequent use. RNA was extracted from each fat depot using Trizol and RNeasy mini kits (Qiagen, Germany). Bioanalyzer (Agilent) analysis was used to perform quality control. cDNA libraries were prepared for samples with RIN > 6. RNA-Sequencing libraries were made using stranded Total RNASeq library prep and libraries were sequenced using Illumina Nextseq500, paired-end read of 75 bp, 50 million reads per library. Illumina adapters of libraries after sequencing were removed using Cutadapt (v1.18)^12^ with default parameters. Sequenced libraries were then aligned to GRCh38 using STAR (v2.42a)^13^ with –twopassMode Basic. RSEM (v1.3.1)^14^ was used for transcript quantification, with rsem-calculate-expression (*--forwad-prob 0.5*).

**Table 1.** Characteristics of TAO and control patients undergoing orbital surgery. Caucasian (C). African American (AA). Uveal Melanoma (UM).

|  | Control S1 | Control S2 | Control S3 | TAO S1 | TAO S2 | TAO S3 |
| --- | --- | --- | --- | --- | --- | --- |
| Age (years) | >70 | 58 | >70 | 37 | 51 | 51 |
| Gender | M | M | F | F | F | M |
| Race | Asian | C | C | C | AA | AA |
| Duration of thyroid disease prior to surgery (approx. mo) | - | - | - | 48 | 13 | 13 |
| Duration of TED prior to surgery (approx. mo) | - | - | - | 18 | 13 | 10 |
| Previous treatment for Graves disease | - | - | - | Armor thyroid | Methimazole | Carbimazole |
| Previous treatment for TED | - | - | - | - | - | Prednisone, Methylprednisolone, Azathioprine |
| Smoking history | >10 years ago | No | No | Former | No | No |
| Exophthalmometry, Hertel (mm) | N/A | N/A | N/A | 21 | 27.5 | 30 |
| Presence of compressive optic neuropathy | N/A | N/A | N/A | No | No | Yes |
| Surgery | Excision of orbital fat prolapse | Excision of orbital fat prolapse | Enucleation (UM) | Orbital decompression | Orbital decompression | Orbital decompression |
| Clinical Activity Score (0-7) | - | - | - | 0 | 1 | 3 |

DESeq2(1.24.0)^15^ was used to analyze the bulk RNA-Seq dataset using the standard pipeline, filtering low counts (<10) and choosing genes with adjusted p-value < 0.05. ClusterProfiler (v3.12.0)^16^ *enrichKEGG* function was used to identify pathways that are enriched in the TAO group.

### *In vitro* differentiation

Orbital fibroblast cell lines were derived from TAO retrobulbar fat as previously described.^17^ Briefly, tissue explants were obtained from patients undergoing surgical decompression for TAO. Patients were inactive, with a CAS less than 4, at the time of surgery.^11^ Explants were placed on the bottom of culture plates and covered with Eagle’s medium containing 10% FBS, antibiotics, and glutamine. They were incubated at 37°C, 5% CO_2_, in a humidified environment. The resulting fibroblast monolayers were passaged serially by gentle treatment with TrypLE. Strains were stored in liquid N_2_ until needed and were used between the 4th and 8th passages.

Orbital fibroblasts were induced to undergo adipogenesis as previously described.^17^ Briefly, fibroblasts between passages 4 and 8 were seeded on plastic tissue culture plates and allowed to proliferate to near-confluence in DMEM containing 10% FBS and antibiotics. The cells were then treated with adipogenic medium consisting of DMEM:F-10 (1:1) supplemented with 3% FBS, 100 nmol/liter insulin, 1 μmol/liter dexamethasone, and for the first week only, 1 μmol/liter rosiglitazone and 0.2 mmol/liter IBMX. Media was changed every other day for the first week, and then twice weekly for the remaining weeks. Cultures were maintained in this medium for 21 days. Control cultures were treated with DMEM:F-10 (1:1) supplemented with 3% FBS and vehicle. Differentiation was observed using a Nikon microscope. Each time point was analyzed in triplicate.

### Quantification of adipocytes

On days 0, 5, 9, and 21, cells were washed with PBS and fixed using 4% paraformaldehyde in PBS for 10 minutes at room temperature. Cells were washed with PBS or stored in PBS at 4°C. To stain adipocytes, cells were washed with 60% isopropanol and stained with freshly prepared 0.3% (w/v) Oil red O at room temperature for 15 minutes. Cells were rinsed with isopropanol and stained lightly with hematoxylin, then rinsed with PBS. Cells were imaged by a blinded study team member using a Keyence BZ-X710 microscope (Keyence, Japan). Four high-power field images were obtained per replicate. Cell counts and lipid vacuole measurements were performed using ImageJ.^18^

### Single-nucleus RNA-Seq

Nuclei from the treated groups were isolated at day 0, 5, 9, and 21 using the methods described in the 10x Genomics Sample Preparation Demonstrated Protocol. Briefly, cells were washed with chilled PBS and lysed in chilled lysis buffer consisting of 10 mM Tris-HCl, 10 mM NaCl, 3 mM MgCl2, and 0.1% Nonidet™ P40 Substitute in nuclease-free water at 4°C. Cells were scraped from the plate bottom and centrifuged at 500 RCF for 5 min at 4°C. Cells were washed twice in nuclei wash and resuspension buffer consisting of PBS with 0.1% BSA and 0.2 U/ul RNase inhibitor. Cells were passed through a 50 um cell strainer and centrifuged at 500 RCF for 5 min at 4°C prior to resuspension in nuclei wash and resuspension buffer. Isolated nuclei were counted manually via hemocytometer with Trypan Blue staining, and nuclei concentration was adjusted following the 10x Genomics guideline. 10x Genomics Chromium Single Cell system (10x Genomics, CA, United States) using V2 chemistry (Day 0, 5, 9) or V3 chemistry (Day 21) per manufacturer’s instructions, generating a total of 7 libraries. Day 5 and Day 21 were run with technical replicates. Libraries were sequenced on Illumina NextSeq500 mid-output or NovaSeq6000 (150 million reads). Sequencing data were first pre-processed through the Cell Ranger pipeline (10x Genomics, Cellranger v5.0.0) with default parameters, using GRCh38-2020-A genome with *include-introns*.

Matrix files generated from the Cellranger run were used for subsequent analysis. Seurat V3^19^ was used to perform downstream analysis, only including cells with more than 500 genes, 1000 UMI, to process control and treated samples separately. Seurat SCTransform function was used to normalize the dataset^20^ and UMAP was used to reduce dimensions derived from the Harmony^21^ output.

In order to identify cells that show adipocytes signatures, key adipocytes-enriched genes were first extracted from GTEx dataset from ascot.cs.jhu.edu^22^, relying on both robustness (NAUC > 20) and specificity (expressed only in both adipocytes dataset or in tissues that are enriched with adipocytes such as mammary tissues). These genes were then used to train the dataset using Garnett^23^ and identified differentiated adipocytes.

RNA velocity^24^ was used to verify *in vitro* adipocyte differentiation trajectory. Kallisto and Bustools^25^ were used to obtain splicing values, and Scanpy^26^ and scVelo^27^ were used to obtain velocity trajectory. Pseudotime analysis was performed using Monocle v3^28^ and genes that are significant (q value < 0.001) along pseudotime trajectory were used to run KEGG analysis as described above. Genes that were involved across KEGG signaling pathways were then used to identify signaling pathways that are specific at different stages of *in vitro* differentiation. SCENIC^29^ was used to calculate regulons controlling gene expressions across adipocytes differentiation stages as previously described.^30,31^ Enriched genes from control and TAO bulk RNA-Seq datasets were superimposed on *in vitro* dataset using Seurat *AddModuleScore* function.

### Data availability

All sequencing data are available on GEO as GSE158464 (bulk RNA-Seq dataset) and GSE174139 (bulk RNA-Seq dataset and snRNA-Seq).

## Results

### Bulk transcriptome analysis of retrobulbar fat from TAO and control patients

Retrobulbar fat was collected from patients undergoing orbital surgery for TAO or other indications (Table 1). To compare gene expression in orbital fat in TAO patients and controls, bulk RNA-Seq was performed (Figure 1a). Gene expression in TAO and control orbital fat segregated using PCA, with control and TAO replicates appearing more similar (Figure 1b). We observed 902 genes that are enriched in control orbital fat, and 964 genes enriched in TAO orbital fat (Figure 1c). Kyoto Encyclopedia of Genes and Genomes (KEGG) pathway analysis of genes enriched in TAO orbital fat revealed numerous signaling pathways that are highly enriched including PI3K-Akt signaling, cAMP signaling, AGE-RAGE signaling, regulation of lipolysis in adipocytes, and thyroid hormone signaling pathway (Figure 1d). Heat map analysis of genes that are differentially expressed (adjusted p-value < 0.05, Table S1) in TAO fat and control fat demonstrated general differences between TAO and control orbital fat, while also identifying differences that correlate with the patient’s clinical activity score (CAS) at the time of surgery (Figure 1e, Table 1). Heat map analysis of individual genes within the above-mentioned pathways demonstrated the degree to which genes such as *THRA* and *IGF1* are enriched in TAO orbital fat (Figure 1f).

**Figure 1.**
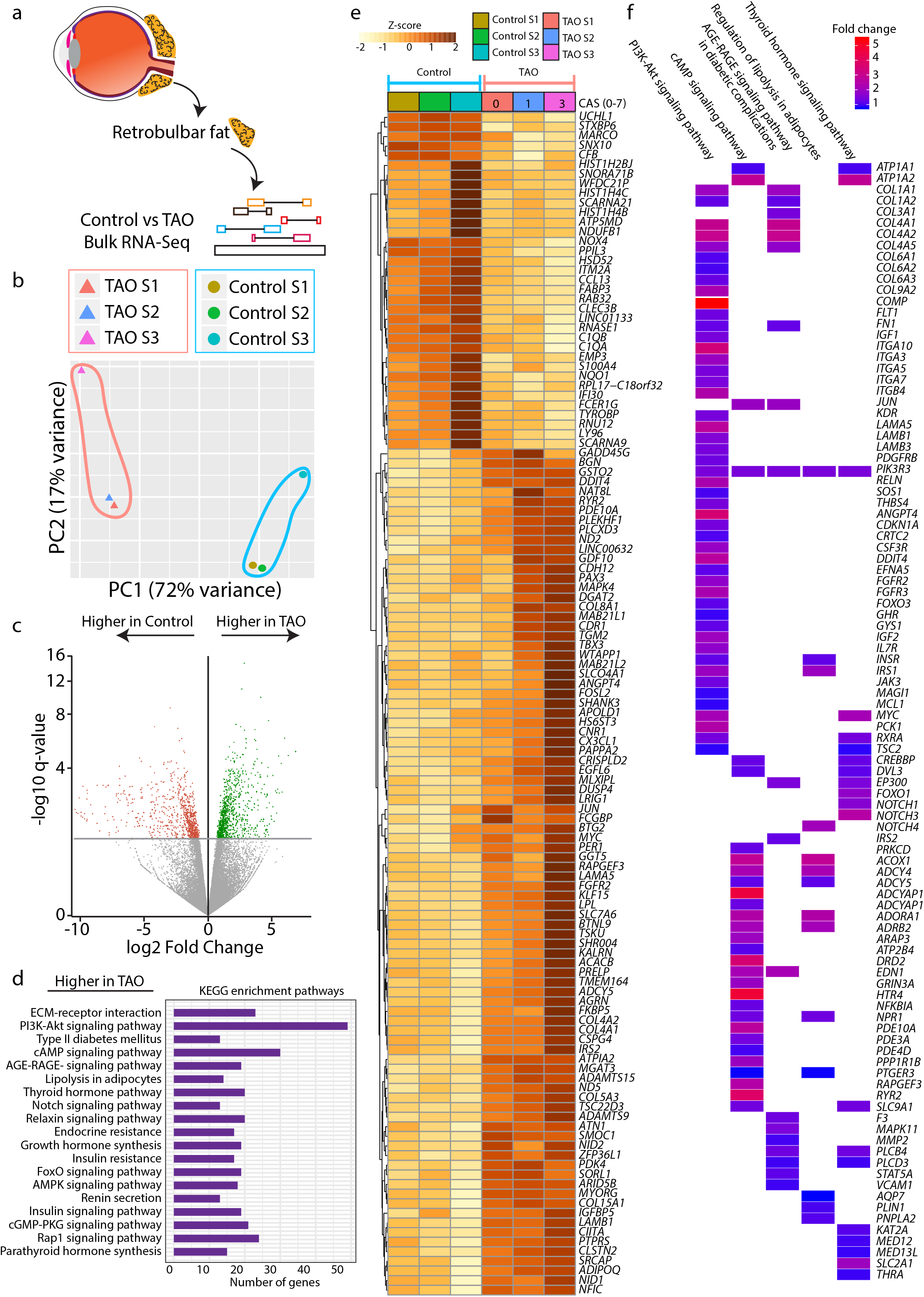
Bulk RNA-Seq analysis comparing retrobulbar fat of control and TAO patients. (a) Schematic diagram showing the overall experiment pipeline. (b) PCA plot showing the distribution of control and TAO replicates. (c) Volcano plot of genes that are higher in control or TAO orbital fat. (d) KEGG pathway analysis showing pathways that are enriched in the TAO group. (e) Heatmap demonstrating gene expression differences between the control and TAO. (f) Top KEGG pathways in 1d and genes that are expressed at a higher level in TAO than in the control group (Fold change). CAS = Clinical activity score.

### snRNA-Seq of orbital fibroblasts undergoing adipogenesis reveals induction of adipocyte-specific markers

Orbital fibroblasts derived from Case #2 were treated to induce adipogenesis using over a 21-day time course to find genes that are induced during adipogenesis which may function critically in orbital fat hyperplasia in TAO. Cells treated with control media did not undergo adipogenesis (Figure 2a,c,e,g). Based on histologic changes noted in cells treated with adipogenic media, we chose to analyze adipogenesis and gene expression at Days 0, 5, 9, and 21 (Figure 2a–h). Adipocyte induction first occurred between day 0-5, which the maximum density of adipocytes noted at day 9 (Figure 2i). Adipocyte maturation, as quantified by the lipid vacuole area, increased most between days 9 and 21 (Figure 2j).

**Figure 2.**
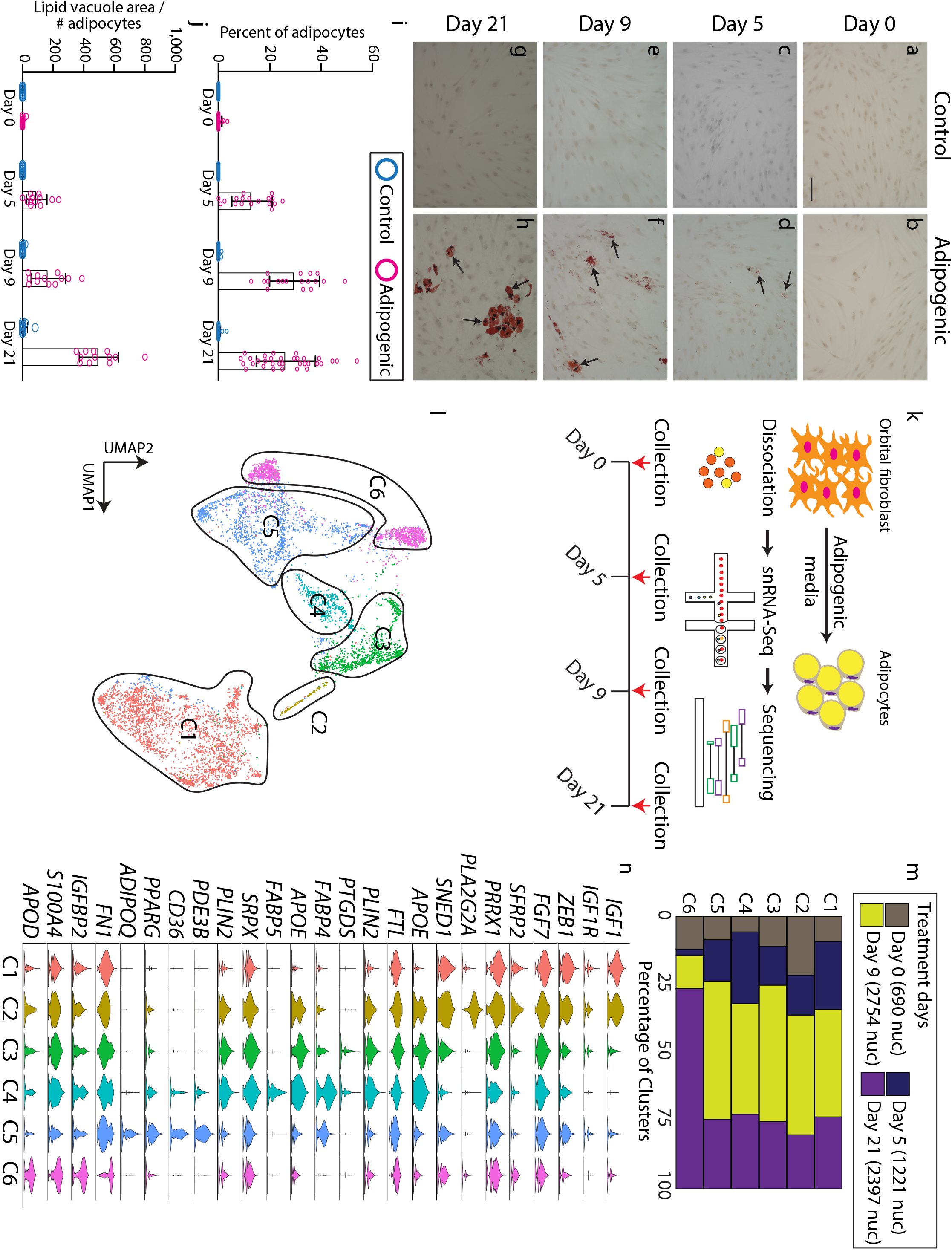
snRNA-Seq shows adipogenesis trajectory *in vitro*. (a-h) Oil Red O staining (arrows) of orbital fibroblasts treated with control (a,c,e,g) and adipogenic (b,d,f,h) medium orbital at day 0 (a,b), day 5 (c,d); day 9 (e,f), and day 21 (g,h). (i) Bar plot showing the percentage of adipocytes seen after treatment with control or adipogenic medium on days 0, 5, 9, and 21. (j) Bar plot showing lipid vacuole area / the number of adipocytes between orbital fibroblasts treated with control or adipogenic medium on days 0, 5, 9, and 21. (k) Schematic diagram showing snRNA-Seq pipeline. (l) UMAP plot showing key 6 clusters of cells undergoing adipogenesis from snRNA-Seq. (m) Bar plot showing the percentage of clusters that are occupied across treatment days. (n) Violin plot showing key cluster markers. Scale bar = 100 μM.

Based on the observed histologic changes, we collected nuclei at days 0, 5, 9, and 21 treatment for analysis by snRNA-Seq (Figure 2k). We identified unique clusters that are enriched from Day 5 onwards and begin to express unique sets of markers that were absent in the rest of the clusters, such as *IGF1* (Figure S1, Table S2). These clusters led to a cluster that expressed high levels of adipocyte signatures including *ADIPOQ* (Figure S1, Table S2). We identified these clusters as cells that are undergoing adipogenesis.

UMAP analysis of differentiating orbital fibroblasts revealed 6 distinct clusters defined by a unique set of genes (Figure 2l). Overall, differentiating cells appeared to progress from cluster 1 (C1) to C6 over time (Figure 2m). Classic adipocyte stem cell markers such as *FABP4*, *APOE*, *FABP5* were enriched in C3-C4, while mature adipocyte markers such as *PPARG* and *ADIPOQ* were enriched in C5, demonstrating that the clusters corresponded to a developmental progression from orbital fibroblast, through adipocyte stem cells, to mature adipocytes (Figure 2n, Table S3). At day 21, a new cell type was present that demonstrated high expression of *APOD*, but had lost expression of other adipogenic markers such as *FABP4/5* but shared expression of many of the other markers expressed in other clusters were represented in cluster C6 (Figure 2n). To identify genes that were induced early in the process of adipogenesis, we focused on genes that were specifically expressed in C1 and C2, which differed from both quiescent, control-treated orbital fibroblasts and adipocyte stem cells. This population showed enrichment of *IGF1*, *IGF1R*, *ZEB1*, *FGF7*, and *SFRP2* expression (Figure 2n). Enrichment of *IGF1* expression within differentiating orbital fibroblasts suggests that paracrine signaling may be involved in the process of adipogenesis.

### Analysis of key pathways involved in orbital adipogenesis *in vitro*

We next focused on identifying key signaling pathways and potential regulatory factors that are differentially expressed during orbital adipogenesis *in vitro*. Using KEGG analysis of the snRNA-Seq data, we observed that differentiating adipocytes express genes involved in multiple signaling pathways, including the PI3-AKT pathway, AGE-RAGE signaling pathway, insulin resistance pathway. These pathways were also enriched in TAO orbital fat compared to control orbital fat, as measured by bulk RNA-Seq analysis (Figure 1c, 3a). Other pathways that are specifically involved in adipogenesis, such as the PPAR signaling pathway, fatty acid biosynthesis and fatty acid elongation pathways (Figure 3a) were also enriched in differentiating orbital fibroblasts.

**Figure 3.**
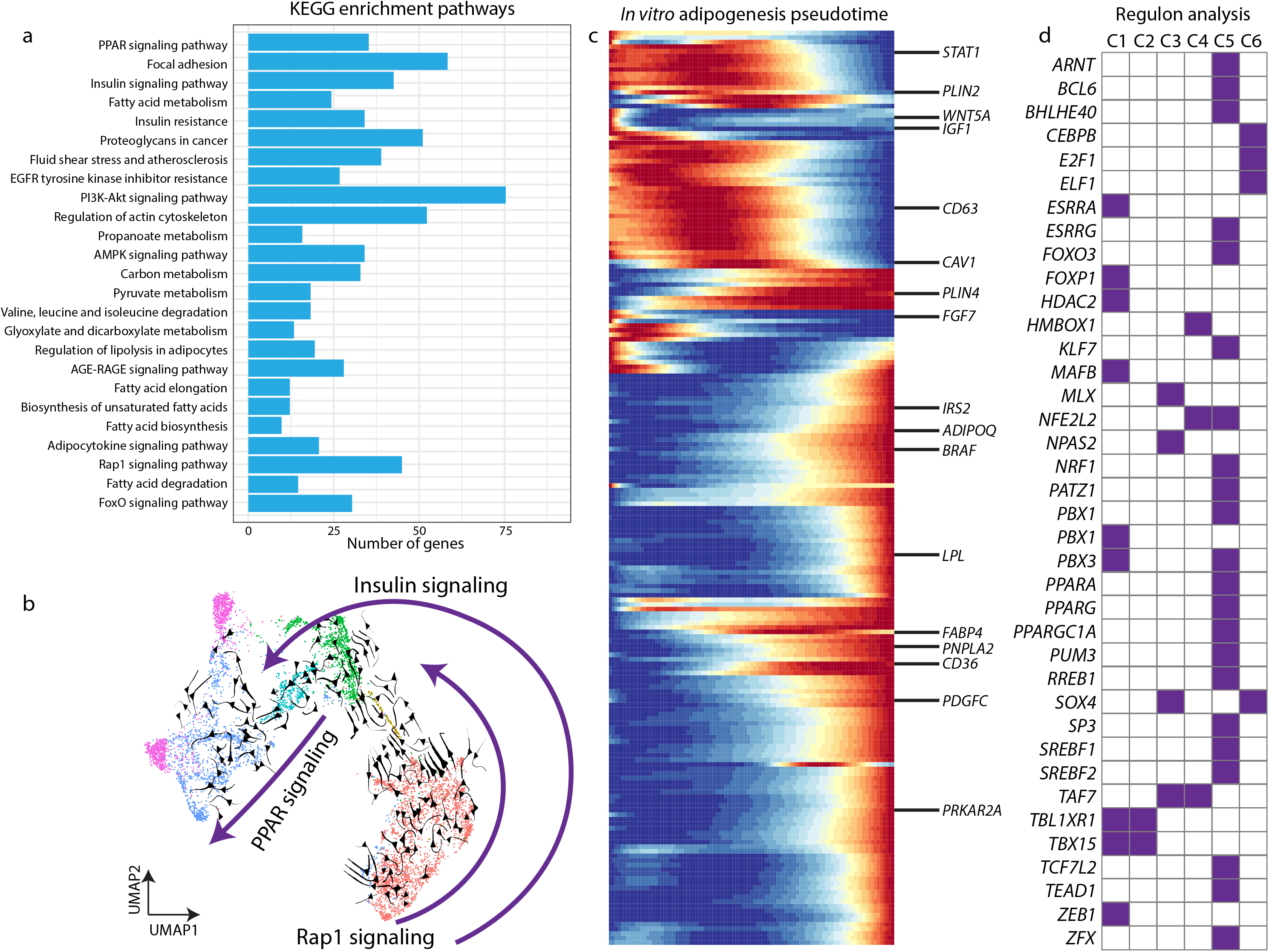
snRNA-Seq reveals adipogenesis signaling pathway. (a) KEGG plot based on pseudotime analysis from snRNA-Seq data in Figure 2. (b) UMAP plot with RNA velocity showing adipocyte differentiation trajectory, and 3 pathways that are specific across differentiation stages. (c) Pseudotime trajectory of adipogenesis and key genes. (d) Regulon analysis demonstrates key transcription factors that might be involved in each step of adipogenesis differentiation.

RNA velocity analysis demonstrated that components of the Rap1 signaling pathway are active early in orbital fibroblasts in C1 (Figure 3b). Components of the insulin signaling pathway were expressed throughout the course of adipocyte differentiation in C1-C5 (Figure 3b), while peroxisome proliferator-activated receptor (PPAR) signaling, which is known to be active late in adipogenesis, was active in C4-C5 (Figure 3b). Pseudotime analysis showed the progression of gene expression through C1-C5 for all differentially-expressed genes (Figure 3c). Genes with particularly strong differential expressions are highlighted, such as *IGF1*, *SFRP2*, and *WNT5A* (Figure 3c, Table S4). Regulon analysis identified transcription factors that are differentially expressed in each cluster (Figure 3d).

### The TAO orbital fat transcriptome resembles that of *in vitro* differentiated adipocytes

In order to determine the similarity between *in vitro* derived orbital adipocytes and orbital fat *in vivo*, we compared gene expression from control and TAO orbital fat, analyzed by bulk RNA-Seq, to gene expression during orbital adipogenesis as assessed by snRNA-Seq (Figure 4a). Control gene modules were found to be expressed most highly in C1 and C6. These modules were most strongly expressed in day 0 cells (C1) and a subset of day 21 cells that do not express mature adipocyte markers (C6) (Figure 4b–d). TAO gene modules were expressed most highly in C5, and at days 9 and 21 (Figure 4e,f). Interestingly, although the orbital fibroblast cell line was derived from a TAO patient, the gene expression pattern resembled control gene modules prior to treatment with adipogenic media (Figure 4c,d), and changes to resemble TAO genes modules following treatment with adipogenic media (Figure 4 f,g).

**Figure 4.**
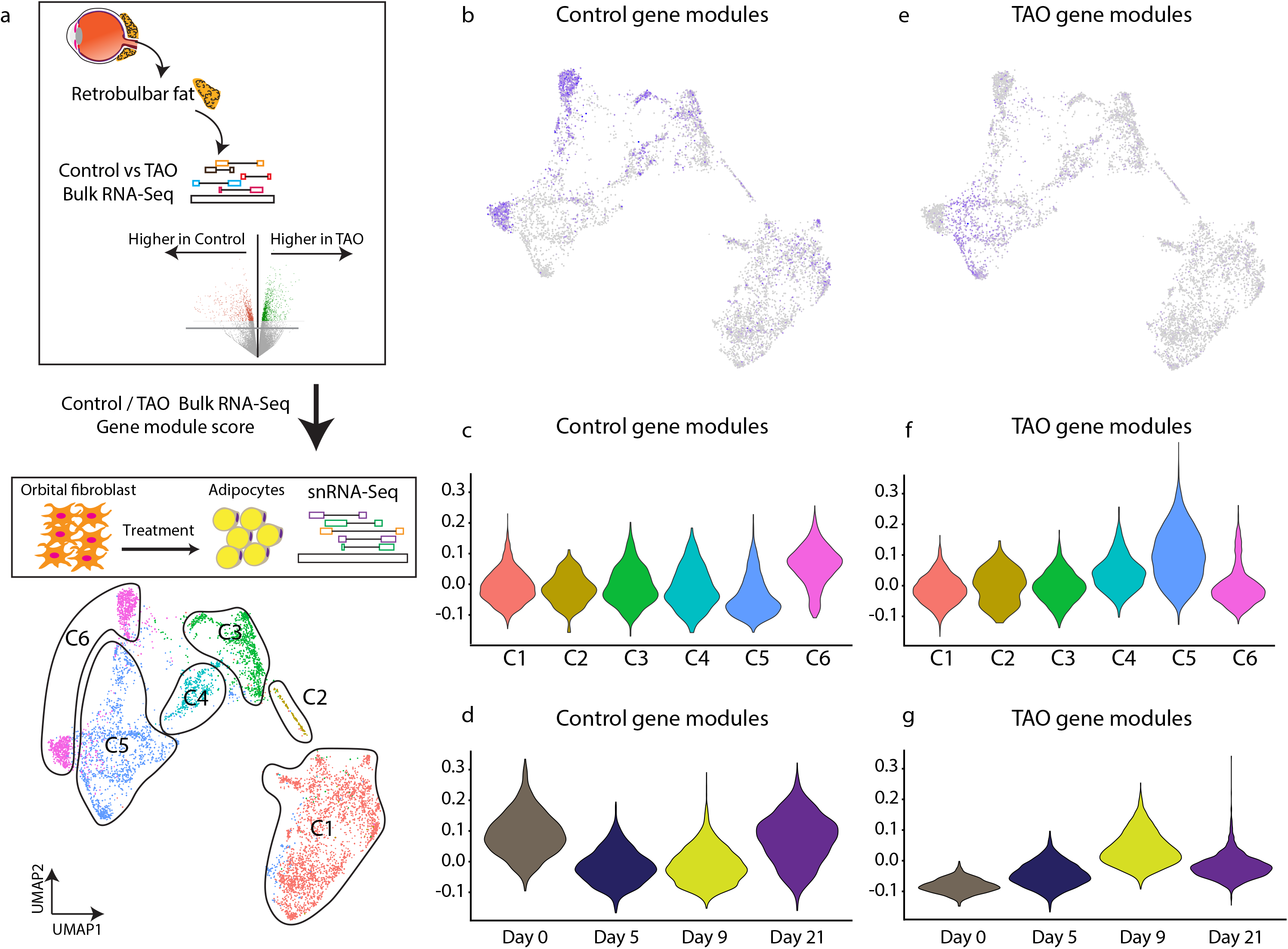
*In vitro* differentiated adipocytes express high levels of TAO tissue-enriched markers. (a) Schematic showing that genes that are higher in control or TAO orbital fat were projected into snRNA-Seq dataset to obtain gene module score. (b-d) Control retrobulbar fat-enriched genes demarcate Cluster 6 in snRNA-Seq dataset (b,d), expressed higher at day 0 (c). (e-g) TAO retrobulbar fat-enriched genes demarcate Cluster 5 adipocytes in snRNA-Seq dataset (e,f), expressed higher at day 0 (g).

## Discussion

In this study, we demonstrate that orbital fibroblasts derived from TAO patients can be used as a model to study adipogenesis *in vitro*. Histologic studies and transcriptome profiling clearly show that, although they are derived from TAO patients, the gene expression profile of undifferentiated orbital fibroblasts closely resembles that of control orbital fat cells, while those treated to undergo adipogenesis display a gene expression profile more similar to that of TAO orbital fat.

We have identified signaling pathways and transcription factors that are upregulated early in orbital adipogenesis, including the IGF-1 signaling pathway, which has previously shown to contribute to TAO pathophysiology by increasing auto-antigen display, cytokine synthesis, and hyaluronan production by orbital fibroblasts.^32,33^ The novel TAO therapy teprotumumab targets the IGF-1 receptor (IGF1-R) to improve proptosis, diplopia, CAS, and quality of life in TAO patients.^34,35^ Though it is thought to exert its exerts via modulation of the immune response, our data raise the possibility that inhibition of the IGF-1R pathway may also impact TAO by reducing adipogenesis in the orbit.

Prior transcriptome profiling of cultured orbital fibroblasts derived from control and TAO patients has demonstrated significantly increased expression of homeobox transcription factors and decreased expression of Wnt signaling pathways components in TAO orbital fibroblasts.^36^ In profiling TAO orbital fibroblasts undergoing adipogenesis, we find that *WNT5A* expression is high early in the course of adipogenesis, but decreases as differentiation occur, whereas homeobox gene expression increases later (Figure 3c,d).

The model that we have used to study gene expression changes during orbital adipogenesis depends on the treatment of orbital fibroblasts with drugs including insulin and rosiglitazone. For example, while cells are treated with insulin throughout the experiment, and the insulin pathway is active throughout the differentiation process, cells are also treated with the PPAR gamma agonist rosiglitazone early (during the first week) in the experiment, but PPAR signaling is active only later in the differentiation process (Figure 3b).

Microarray comparisons of orbital fat from TAO and control patients have demonstrated increased expression in Wnt signaling pathway genes such as SFRPs, IGF-1 pathway genes, and adipogenic genes, consistent with our bulk RNA-Seq results.^37,38^ In our heat map analysis of TAO-enriched genes, we note differences in the level of gene expression that correlate with CAS. Case 1, with a CAS of 0, has lower levels of TAO-enriched genes than Case 2, with a CAS of 1, which in turn has lower levels of TAO-enriched genes than Case 3, with CAS of 3 (Figure 1e). However, even in the least severely affected patient, TAO-enriched genes are more highly expressed than in controls (Figure 1e).

Other studies have compared RNA-Seq data of orbital fat from patients with active TAO compared to blepharoplasty fat and observed stronger upregulation of inflammatory signaling pathways, which is consistent with the higher CAS in their patients compared to those profiled in our study.^39^ Conversely in our patients, pathways including insulin and thyroid signaling, as well as adipogenic pathways, were more prominent, likely reflective of the patients being in earlier stages of the disease. In addition, we used retrobulbar fat as our control orbital adipose sample, rather than blepharoplasty fat, which may account for some of the other observed differences in gene expression. A potential confounding factor in comparing these data sets may be that in our analysis, we have excluded all genes with low counts due to low expression levels. Nonetheless, we also found large changes in Wnt signaling such as (*DKK2*, *NKD1*) and other pathways (*HOX3D*) in TAO orbital fat compared to control orbital fat in post-hoc analysis (Table S1), although these genes were expressed at low levels.

We further have identified factors that have not been previously implicated in the biology of orbital fibroblasts, including cell-autonomous pathways such as Rap1 signaling, which have been shown to drive adipogenesis in human bone mesenchymal stem cells.^40,41^ In addition, transcription factors that drive adipogenesis are of particularly interest in understanding the regulation of orbital adipogenesis. ZEB1 has been identified as a critical mediator of adipogenesis in mouse pre-adipocytes and is induced by IGF-1R activation in prostate cancer cell lines.^42,43^ Finally, the use of *in vitro* analysis of differentiating cultured orbital fibroblasts to identify pathways that regulate orbital adipogenesis, and that are enriched in TAO orbital fat, opens the possibility of using high-throughput screening to identify drug and gene-based approaches to modulating pathological adipogenesis that may ultimately be useful in treating TAO-related disorders.

## Acknowledgements

This work was supported by an unrestricted departmental grant to the Wilmer Eye Institute from Research to Prevent Blindness. DWK is supported by the Maryland Stem Cell Research Fund (2019-MSCRFF-5124). SB is supported by NIH Grant R01EY020560. FR is supported by the National Eye Institute of the National Institutes of Health under award number K08EY027093. We thank the Transcriptomics and Deep Sequencing Core (Johns Hopkins) for sequencing of snRNA-Seq libraries. We also thank Lizhi Jiang for technical assistance.

---

**Figure S1. snRNA-Seq of the overall dataset.** (a,b) UMAP plot showing the distribution of clusters across the entire dataset (a) and distribution across each treatment age or replicates (Day 9 and Day 21) (b). Red lines in panels a and b indicate cells are undergoing adipogenesis. (c) Gene expression showing unique genes that are enriched in cells that are undergoing adipogenesis.

**Table S1.** Significant genes from bulk RNA-Seq of control and TAO retrobulbar orbital fat in Figure 1.

**Table S2.** Cluster markers of Figure S1a.

**Table S3.** Cluster markers of Figure 2l.

**Table S4.** Pseudotime genes of Figure 3c.

