## Supplementary figures and images for "Transcriptomic profiling of human orbital fat and differentiating orbital fibroblasts"

### Figure S1

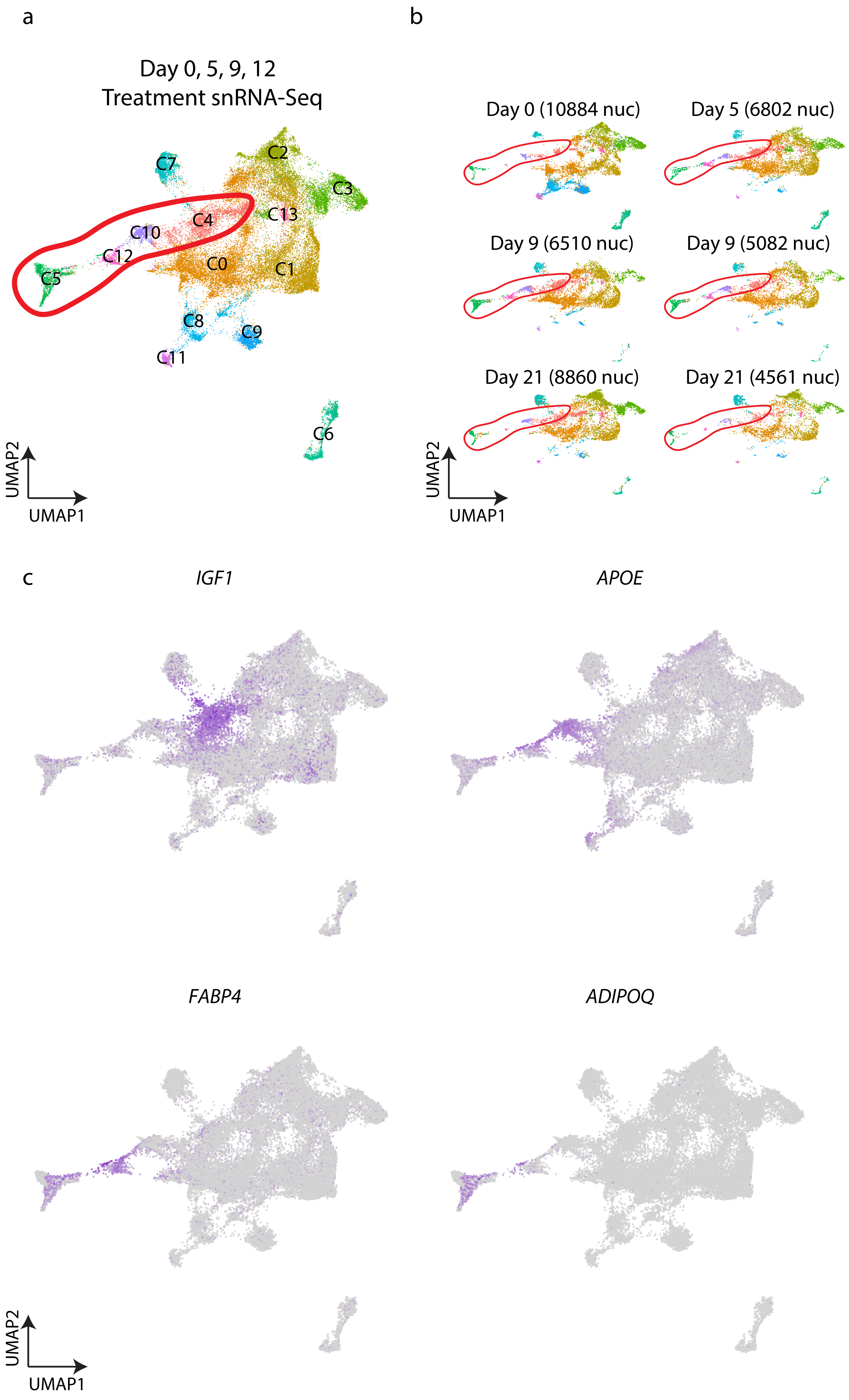
